# The future burden of congenital Toxoplasmosis in Africa under demographic and climate change

**DOI:** 10.1101/2025.09.04.674311

**Authors:** F. Rasambainarivo, I.G. Nilsson, D. Cheeks, Wenchang Yang, C.J.E. Metcalf

## Abstract

The impact of climate change on environmental pathogens is a question whose importance will amplify in coming years. The protozoan parasite and global zoonosis *Toxoplasma gondii* is one such: empirical evidence indicates that oocyst survival is reduced at high temperatures. Paradoxically, a decline in incidence of *T. gondii* infections could amplify the burden of this disease, as the most damaging outcome occurs subsequent to first infection during pregnancy, and reductions in the incidence of the infection will increase the average age of first infection. We blend models of infection dynamics rooted in occurrence across the African continent with models of human demography to bound expectations for the future burden of this pathogen, accounting for the effects of changing temperatures. We discuss targeting efforts and approaches for mitigation.

## Introduction

Toxoplasmosis, caused by the protozoan parasite, *Toxoplasma gondii*, is a worldwide zoonosis (Figure 1) and a leading cause of severe foodborne infection [1,2]. Infection poses risks to vulnerable populations, notably pregnant women as the parasite can be transmitted to the fetus causing congenital defects including occulitis, encephalopathy, fetal death and stillbirth, a set of outcomes termed ‘congenital toxoplasmosis’ [3]. It has been estimated that congenital toxoplasmosis leads to more than 800,000 disease-adjusted life years (DALYs)/year globally [2], particularly in South America and sub-Saharan Africa [4,5].

**Figure 1:**
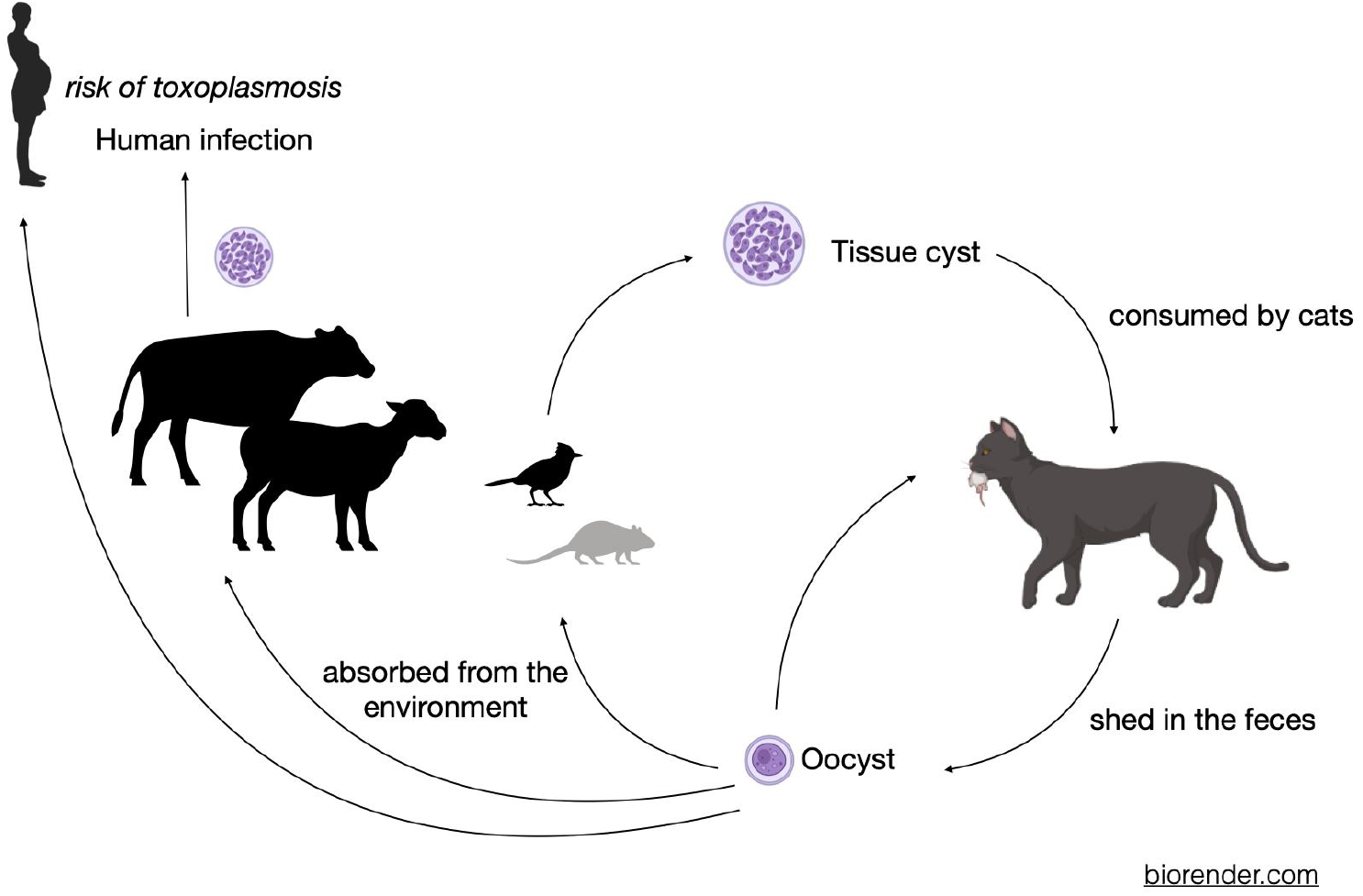
Life cycle of *T. gondii*: Members of the Felidae family (including domestic and wild cats) are the only definitive hosts of *T. gondii*. Cats and wild felids are infected by consuming the tissues of an infected intermediate host, ingesting oocysts in the environment, or through congenital transmission (not shown). Oocysts are then shed in the environment through the cats’ feces and sporulate 1 to 5 days later, becoming infective to other intermediate or definitive hosts. Humans and other intermediate hosts can also become infected by consuming food or water contaminated with oocysts as well as the consumption of infected intermediate hosts.

The large majority of cases of congenital toxoplasmosis occur in women experiencing their first infection during pregnancy [6], although cases in women following infection with a different serotype, or resurgence in immunosuppressed mothers have also been reported [7]. This type of risk profile, where the highest burden is concentrated in women experiencing a primary infection during pregnancy can result in counterintuitive effects of changes in the force of infection, defined as the probability that susceptible individuals become infected. As described for rubella [8], if the force of infection is very high, most individuals will be infected for the first time early in life, and thus will not be at risk during pregnancy; if the force of infection is very low, very few individuals will be infected at all, and thus the burden will be low. Intermediate levels of the force of infection may thus convey the greatest burden within a population (Figure 2).

**Figure 2:**
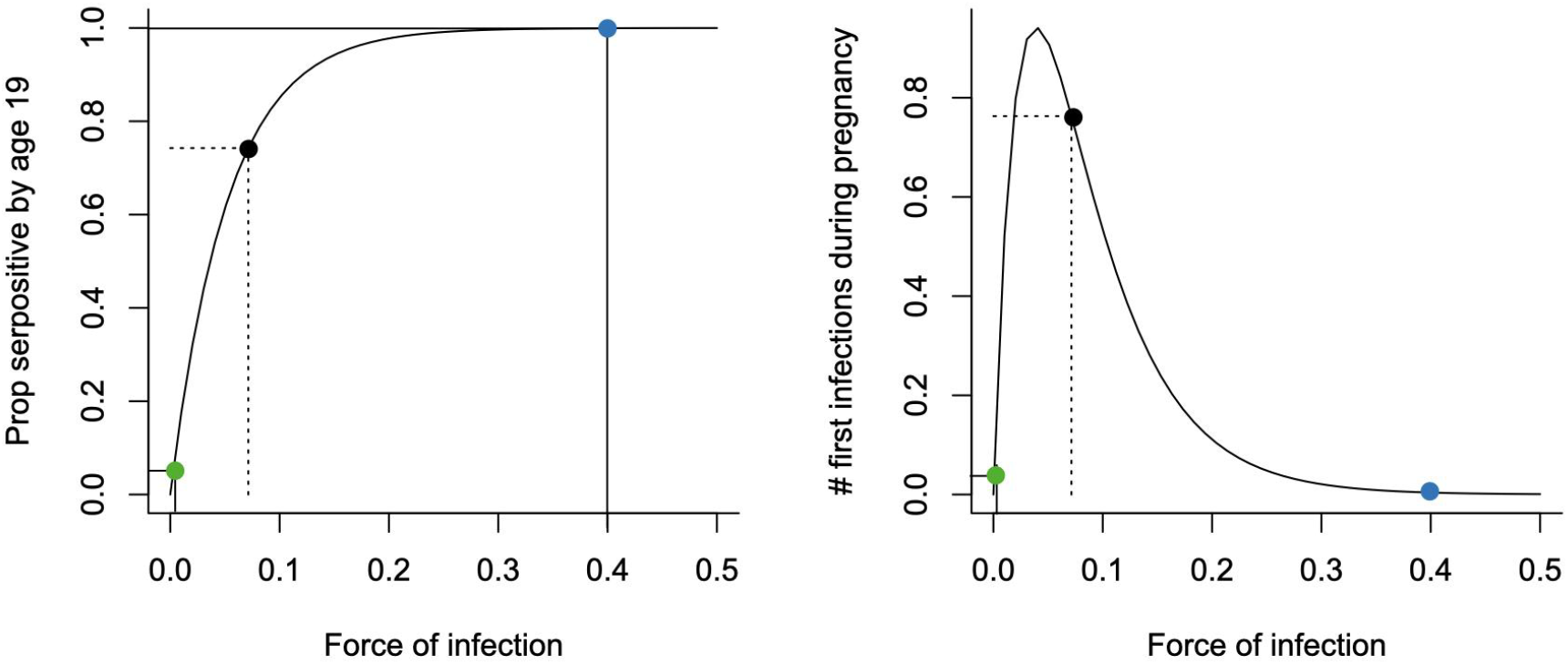
Schematic indicating the paradoxical effect relating the force of infection (FOI) and burden of congenital toxoplasmosis. For individuals living in a location with a high force of infection (blue) almost everyone will already have been infected by age 19, which will mean that very few individuals will be at risk of acquiring a new infection during pregnancy; conversely for individuals living in location with a very low force of infection (green), very few individuals will be infected at all, so although many individuals remain susceptible by age 19, they are at little risk of acquiring the infection. Individuals living in settings with an intermediate force of infection (black) are potentially at the highest risk; thus, paradoxically, a decline in the FOI can result in an increase in burden depending on details of distribution of fertility and how this also changes.

Age seroprevalence studies can be used to estimate the force of infection, revealing patterns of risk across different settings [9]. Age seroprevalence for *T. gondii* in human populations is likely to be context specific, as the life cycle of this protozoan pathogen is mediated by behavioral and social features, such as food preparation methods, but also the environment (Figure 1). Oocysts are shed in the feces of cats, and their survival in the environment is reduced at high temperatures [10]. This latter point was our focus in this paper: the age specific manifestation of the main burden of this infection means that the effects of climate change on oocyst survival might result in large shifts in the impact on population health. However, other dependencies including on local demography mean that the mapping is unlikely to be straightforward.

Here, we combine seroprevalence studies across Africa to estimate the force of infection of this pathogen in different settings through time, and estimate how this is modified by temperature and the human development index, used as a proxy for a suite of hygiene features likely to reduce the risk of toxoplasmosis infection. These estimates of the force of infection and its dependencies allow us to estimate the risk of primary infection across age and combine this with current and future estimates of fertility to evaluate the changing burden of congenital toxoplasmosis, comparing the effects of demographic change (specifically, age structured and fertility profiles in 2023, 2050 and 2100), and temperature change, here evaluating the extreme scenario of a three degree increase in temperature. Our analysis provides a framework to anticipate future disease burden and to identify priority regions for targeted interventions under climate and demographic change.

## Methods

### Serology data

Seroprevalence studies categorize individuals as seronegative, i.e., having never been infected by *T. gondii* (*Y*_*i*_ = 0), and seropositive, i.e., chronically infected (*Y*_*i*_ = 1). Totals by age class are reported across the literature. We leveraged a meta-analysis of seroprevalence across pregnant women [9], expanding the dataset by identifying additional studies through snowballing. Eligible studies reported seroprevalence in pregnant women and provided sufficient information on age groups, sample sizes, and prevalence estimates by age. We included studies that used any diagnostic method for detecting *T. gondii* antibodies (Figure S1: flowchart), restricting the search to those conducted in Africa since 1979 when detailed temperature data became available. A total of 87 articles documenting the seroprevalence of *T. gondii* in 83 919 pregnant women from a total of 26 countries in years from 1979 to 2023 were reviewed and analyzed in this study. Data for each article were extracted using a previously prepared checklist including Name of first author, Year of publication, year of sampling, sample size, age range, number of seropositives (or percentage of seropositives) for each age class (Supplementary Table S1). The full range of the data is shown in Figure 3A and Supplementary Figure S3, and comprises 402 data-points across 25 countries.

**Figure 3:**
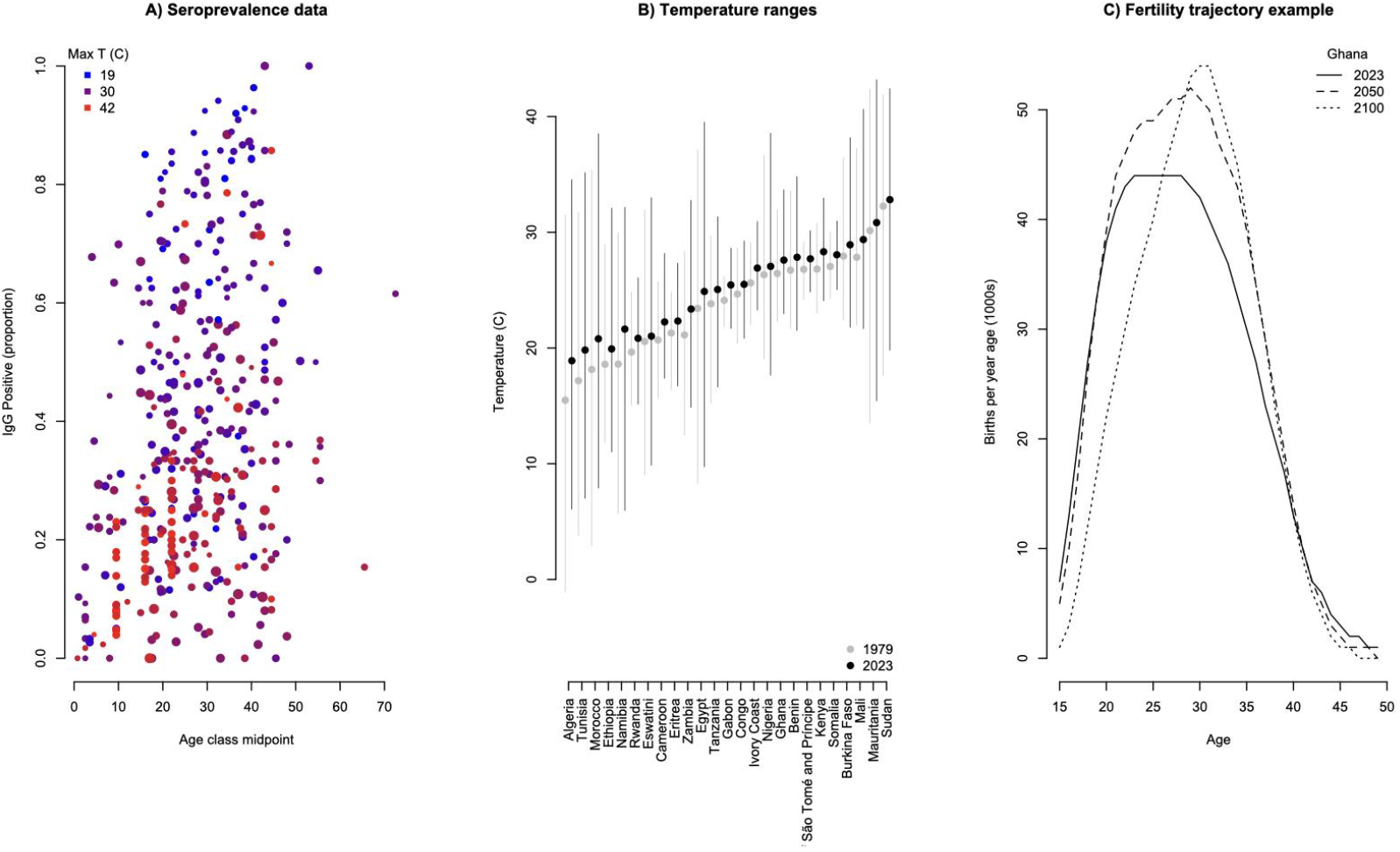
Data used in the analysis including A) Seroprevalence extracted from the literature (coloured by average country temperature at time of study); B) 1979 and 2023 temperature range across the year from the ERA5 reanalysis [11], and C) Fertility profiles from the United Nations Population Division [32], indicating the trajectory of Ghana for illustration.

### Other covariates

Previous work indicates that climatic variables including temperature might alter pathogen ecology, with, in particular, lower survival of oocysts at higher temperatures [10], see Supplementary Figure S2. For each study available, we therefore obtained minimum and maximum temperature using the nearest urban center as described and using the daily ground temperature over the years 1979-2024 from the fifth generation European Centre for Medium-Range Weather Forecasts atmospheric reanalysis (ERA5) of the global climate [11] (Figure 3B). ERA5 combines both in-situ/satellite observations and weather forecast model simulation information to create the best estimate of global-coverage climate information on regular longitude/latitude grids (with an approximate horizontal resolution of 30km). It is a popular dataset used in climate studies and often treated as “observations” when compared to climate model simulations. Ground temperatures at locations of interest in this study, given their information of longitude and latitude, are obtained by bilinear interpolation from the globally gridded data. For serology data preceding the first available temperatures measurements (Mauritania, 1973) we took values representing 1979. Excluding these measurements did not qualitatively alter results

The complex ecology of this pathogen (Figure 1) indicates that different patterns of sanitation and cultural practices (food preparation) can modulate the risk of human infection. Addressing this in detail would require much finer anthropological and social data than currently available; however, larger scale proxies may capture some of the key details. For each country/year combination, we extracted the Human Development Index from OurWorldInData, a metric that blends education, health and economic indexes [12]. To capture additional cultural and other contextual variation, we fit ‘country’ as a random effect in what follows.

### Estimating the force of infection

Data indicating seroprevalence by age opens the way to estimating the force of infection, or rate at which susceptible individuals become infected, defined as λ. In the simplest analysis, the probability of evading infection up until time of sampling t is *S*(*t*) = *exp*(− λ*t*), and the probability of having become infected by time *t* is the complement of this *F*(*t*) = 1 − *exp*(− λ*t*). It follows that *Y*_*i*_ ∼*Bernouilli*(*F*(*t*_*i*_)) and we can thus use a generalized linear model framework to estimate the force of infection [13]. To formally account for covariates modulating the force of infection, we use the complementary log-log link:

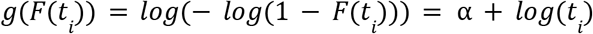

where α = *log*(λ) ; a relationship that can be fit using the glm function in R, setting log age as an offset. To reflect the fact that the force of infection might be modulated by temperature [10] as well as social context features, we assume that

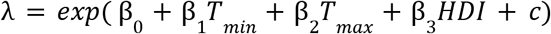

where β_0_ is an overall intercept, β_1_ and β_2_ are slopes on minimum and maximum annual temperatures respectively, β_3_ is a slope on the human development index and *c* is drawn from a distribution centered on zero capturing the range of country specific effects. We fit this model using the lmer package in R where the full link function is defined by:

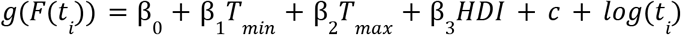

and obtain estimates and standard errors of the full set of parameters.

### Impact on burden of congenital toxoplasmosis

Increasing the force of infection, λ, or the rate at which susceptible individuals become infected increases the average age of first infection, *A*, with *A* = 1/λ. Changes in global demography shift the age at which women are having children (Figure 3C), via both changes in the age distribution of the population, and changes in the age at which women choose to have children. The burden of congenital toxoplasmosis will be determined by the intersection between these two processes (schematically depicted in Figure 2), with details of the magnitude of effects contingent on the details of the associated distributions. To capture this, we use estimates and forecasts provided by the UN Population Division, focussing on ‘Births by Age of Mother’ [14], which accounts for both shifts in age structure, and changing patterns of fertility over age. We do not provide estimates of absolute burden, but rather focus on the relative burden, *r* (reflecting now vs. future scenarios), in order to broadly generalize across the nuances of timing of infection over the course of pregnancy relative to circulating serotypes, differential virulence of different strains in different regions [15], etc. The probability that an individual is infected at age *a* is defined by:

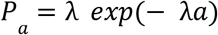

where the second term *exp*(− λ*a*) captures the probability of remaining susceptible up to age a, and the first term the risk of infection at that age. Thus, the relative burden for each country is defined by:

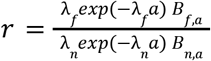

where *B*_*f*,*a*_ is the future number of children born to women of age *a* (either in 2050 or 2100, using the ‘future medium variant’ provided by the UN Population Division) and *B*_*n*,*a*_ is the current (or most recent estimate, i.e., 2023) number of children born to women of age *a*; and λ_*f*_ is the predicted future hazard of infection, λ_*n*_ is the current hazard of infection, based on the most recent estimates of HDI (2022) and temperature (2024).

For λ_*f*_, we explore scenarios where the future climate remains constant (such that only the demography changes via *B*_*f*,*a*_ and *B*_*n*,*a*_, and we retain minimum and maximum temperatures reflecting 2024), or that under future climates, the minimum 2024 temperature is retained, but the maximum temperature increases by 3 degrees celsius, vice versa, or both. In all scenarios, to isolate the effects of climate and demography, we assume that the country-specific HDI is constant at levels reported for 2022.

## Results

The fitted model explains 52% of the variation in seroprevalence and indicates that the force of infection declines with the HDI (Figure 4A) and maximum temperature (Figure 4B) and increases with minimum temperature (Figure 4C), also varying across countries (Figure 4D). The width of age bins across the different studies is highly variable; to verify that our results were not sensitive to this, we estimated relationships using both the midpoint of age bins, and the upper age of age bins; finding largely consistent results.

**Figure 4:**
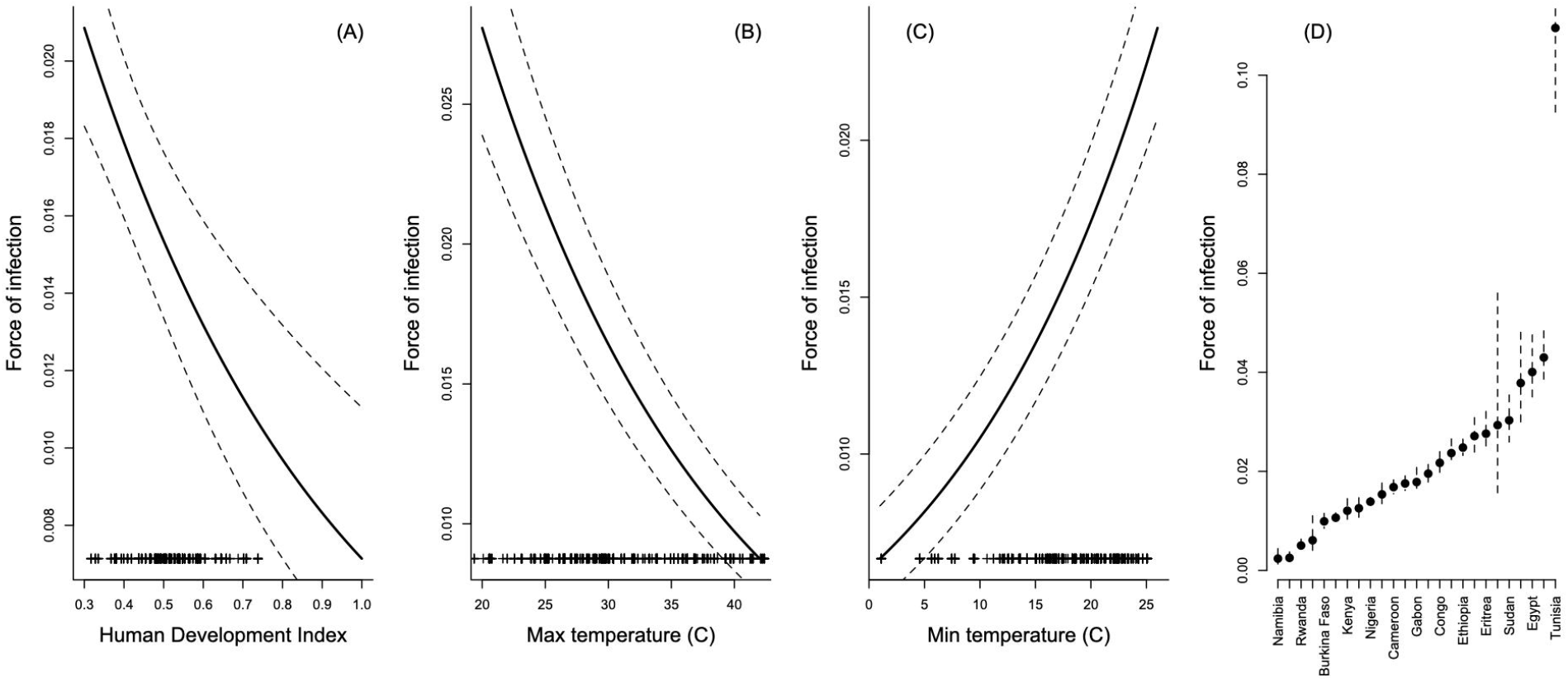
Fitted estimates for the force of infection. (y axis) with mean (solid line) and standard error (dashed line) across variables on the x axis; other variables are set to their average within the data-set. Tickmarks on the x axis indicate the location of the data. Results indicate (A) declines with the human development index (HDI, x axis), a proxy for changes in access to sanitation, etc; (B) declines with maximum temperature (x axis), in line with estimated negative effects of increased temperature on oocyst survival (see Supplementary Figure S2); (C) increases with minimum temperature (x axis), suggesting a lower bound on oocyst survival, as observed in many biological systems; and (D) country level effects capturing residual variation (see Table 1). The full model explains 52% of the deviance.

Combining these results with estimates of current and future births across age [16], assuming a constant HDI (implying broadly invariant sanitation and dietary features in populations), and temperatures either remaining constant or increasing by 3 degrees celsius (the upper limit of expectations), we find a vast diversity of outcomes, ranging from halving to near doubling of the burden of congenital toxoplasmosis (Figure 5, left). Demographic effects largely dominate (the distance between the filled and hollow black points is greater than their distance to coloured points, Figure 5). An increase in maximum temperature in general decreases the burden of congenital toxoplasmosis, while effects of increases in the minimum temperature or both minimum and maximum temperature are variable. The direction of effects resulting from demographic changes are variable: some countries experience an increase in burden (e.g., Tanzania, Ivory Coast) and some a decrease (e.g., Morocco, Tunisia).

**Figure 5:**
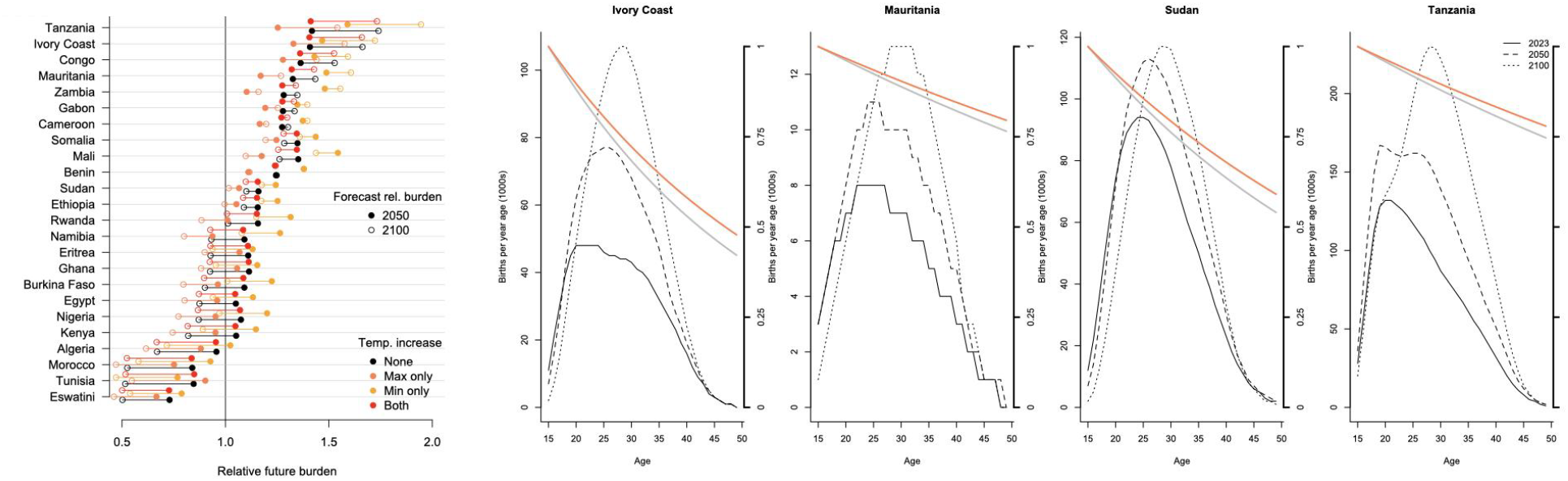
Relative future burden. calculated as the total number of children born to a mother experiencing their first infection during pregnancy and thus at risk of congenital toxoplasmosis in 2050 (filled points) and 2100 (hollow points) relative to 2023, indicating effects at contemporary temperatures (black), and under increases of 3 degrees of the maximum (orange), or minimum (coral) or both (red). Countries that are examples of extreme demographic and temperature effects are depicted, showing fertility over age (line types, legend, left hand y axis) and the proportion of the population experiencing their first infection at each age in current temperatures (grey) and at three degrees higher for both minimum and maximum temperatures (red, right hand y axis). Note that no fertility estimates are available for São Tomé and Príncipe.

To unpack this complexity, we fitted a linear model to log projected burden, with explanatory variables including the type of temperature change (no change, increase maximum, increase minimum, increase both), the year of demographic projections (current, 2050, 2100), the average country HDI, and the interaction between these latter two, leveraging covariation between demographic patterns and HDI to unpack the effect of different time-horizons of demographic change. Country was also fitted as a random effect. Fitted estimates (Table 2) indicate that, on average, increases in the maximum temperature reduce the burden; increases in the minimum temperature increase it; and the burden increases with year more in countries with a higher HDI, suggesting that their demographic structure translates into a concentration of risk in women of childbearing age in the future. Lower HDI countries may thus reflect demographic scenarios where both high force of infection and early age of fertility combine to keep risk low even in the future.

**Table 1:** Parameter estimates for the model fitted to force of infection. (Figure 4) are: an intercept of -2.284 (standard error 0.348), a coefficient on HDI of -1.500 (0.351), a coefficient on maximum temperature of -0.067 (0.003) and a coefficient on minimum temperature of 0.060 (0.004); country specific estimates are provided below.

| Country | Estimate | Standard Error |
| --- | --- | --- |
| Algeria | -3.275 | 0.122 |
| Benin | -4.378 | 0.079 |
| Burkina Faso | -4.615 | 0.077 |
| Cameroon | -4.085 | 0.043 |
| Congo | -3.830 | 0.051 |
| Democratic Republic of São Tomé and Príncipe | -3.147 | 0.060 |
| Egypt | -3.217 | 0.087 |
| Eritrea | -3.590 | 0.077 |
| Eswatini | -3.529 | 0.330 |
| Ethiopia | -3.697 | 0.034 |
| Gabon | -4.026 | 0.078 |
| Ghana | -3.743 | 0.058 |
| Côte d'Ivoire (Ivory Coast) | -3.935 | 0.046 |
| Kenya | -4.419 | 0.094 |
| Mali | -5.102 | 0.303 |
| Mauritania | -4.043 | 0.043 |
| Morocco | -3.607 | 0.066 |
| Namibia | -6.040 | 0.313 |
| Nigeria | -4.278 | 0.029 |
| Rwanda | -5.293 | 0.116 |
| Somalia | -4.176 | 0.071 |
| Sudan | -3.496 | 0.080 |
| Tanzania | -4.543 | 0.044 |
| Tunisia | -2.211 | 0.088 |
| Zambia | -5.979 | 0.199 |

**Table 2:** Parameters for a model synthesizing effects on projected log burden. Temperature change is fitted as a factor with four levels including ‘no change’ ‘change of maximum only’ ‘change of minimum only’ and ‘change of both’; where change describes an increase of 3 degrees celsius, reflecting the extremes; n=288. Country was also fitted as a random effect; the model explained >99% of the variation (unsurprisingly, as it is fitted to model predictions).

| Parameter | Estimate | Standard Error | P-value |
| --- | --- | --- | --- |
| Intercept (no temperature change) | 90.562 | 11.227 | <0.001 |
| Change of max temperature | -0.092 | 0.025 | <0.001 |
| Change of minimum temperature | 0.079 | 0.025 | <0.001 |
| Change of both | -0.003 | 0.025 | >0.5 |
| Year | -0.039 | 0.005 | <0.001 |
| HDI | -159.738 | 20.156 | <0.001 |
| Year x HDI | 0.070 | 0.009 | <0.001 |

## Discussion

Global change will modulate the future burden of toxoplasmosis via multiple channels [17], including changes in cultural and sanitation practices, changes in temperature affecting pathogen survival, and demographic changes in human populations. Here, we leveraged existing age-serology patterns from across the African continent to provide quantitative estimates of how the broad human context (evaluated via the human development index) and maximum and minimum temperature modulate risk of infection. Our results align with expectations: risk declines with HDI and maximum temperature - and also enable us to explore future burden under projections of demographic change across a range of temperature changes.

Our statistical analysis has a number of limitations. We did not fit variation in risk of infection with age, as the data would not allow it. The true relationship with climatic variables is likely to be more nuanced than dependence only on minimum and maximum temperature (e.g., effects of rainfall have been detected [18]). We also did not have data available to address potentially different burdens associated with variation across infecting genotypes in severity and burden of CT [19], or risk associated with infection in mothers immune to a typical genotype challenged by an atypical genotype during pregnancy [20]; although our focus on relative country comparisons should control for this as long as there are no major future changes in global genotype circulation, or infections associated with travel or transport of consumables. Projections also neglect the role of other potentially important drivers of change, such as an increase in consumption of meat, or improvements in hygiene (noting that the two might counterweight each other). Nevertheless it provides a first foundation for considering the future burden of this global and important pathogen.

Temporal declines in *T. gondii* seroprevalence have been observed in several human populations, potentially indicating a reduction in congenital toxoplasmosis (CT) incidence in some settings [21]. The drivers of this decline are likely multifactorial and may include improvements in sanitation and hygiene, better food safety practices, and cat population management. However, our synthesis of projected burdens under future demographic and climatic scenarios suggests a complex, and at times counterintuitive, risk landscape for congenital toxoplasmosis.

This dynamic set of possible outcomes for CT parallels projections for congenital rubella syndrome (CRS) associated with introduction of vaccination [22,23]. Pre-vaccination, most rubella infections occur during childhood, and are relatively mild. Introduction of the vaccine at scales insufficient to achieve herd immunity, has the potential to shift susceptibility into reproductive ages, increasing incidence of CRS. This ‘paradoxical’ outcome where partial control measures can inadvertently heighten the risk to fetuses is referred to as a peak shift dynamic [24]. A similar dynamic could emerge for CT: without appropriate adaptation of public health strategies, declining *T. gondii* exposure may increase susceptibility among women of reproductive age, particularly in populations undergoing rapid urbanization or socioeconomic transition, or rapid increases in temperature. Anticipating this shift will require proactive targeting of interventions. In particular, screening and education programs for at-risk populations (women of childbearing age) should be prioritized in regions where burden is projected to rise. At present, prenatal serological screening remains the only diagnostic tool capable of identifying all potential cases of congenital toxoplasmosis (CT) in time to provide appropriate diagnosis, treatment, and follow-up [25,26]. This approach should be complemented by strengthening and ensuring access to diagnostic capabilities and potentially prophylactic measures in at-risk regions.

While no human vaccine is yet available, development efforts are ongoing [27,28] and the eventual deployment of any vaccine would need to be guided by robust cost–benefit analyses to avoid unintended epidemiological consequences. Lessons from other systems highlight the importance of these economic considerations as they may shape the uptake and effectiveness in a future human *T. gondii* vaccination program. A One Health approach that coordinates across human, animal, and environmental health sectors will be critical to manage transmission risk holistically [29,30]. Such strategies could include targeted vaccination of cats to reduce environmental contamination by limiting oocyst shedding [31].

Finally, this work contributes to a growing body of evidence that climate change does not uniformly reduce or exacerbate disease risks, but instead reshapes the distribution and nature of those risks across populations, and that future human demography remains a key determinant of disease burden.

## Notes

### Competing Interest Statement

The authors have declared no competing interest.

