## Supplementary Figure for "The future burden of congenital Toxoplasmosis in Africa under demographic and climate change"

**Supplementary Materials**

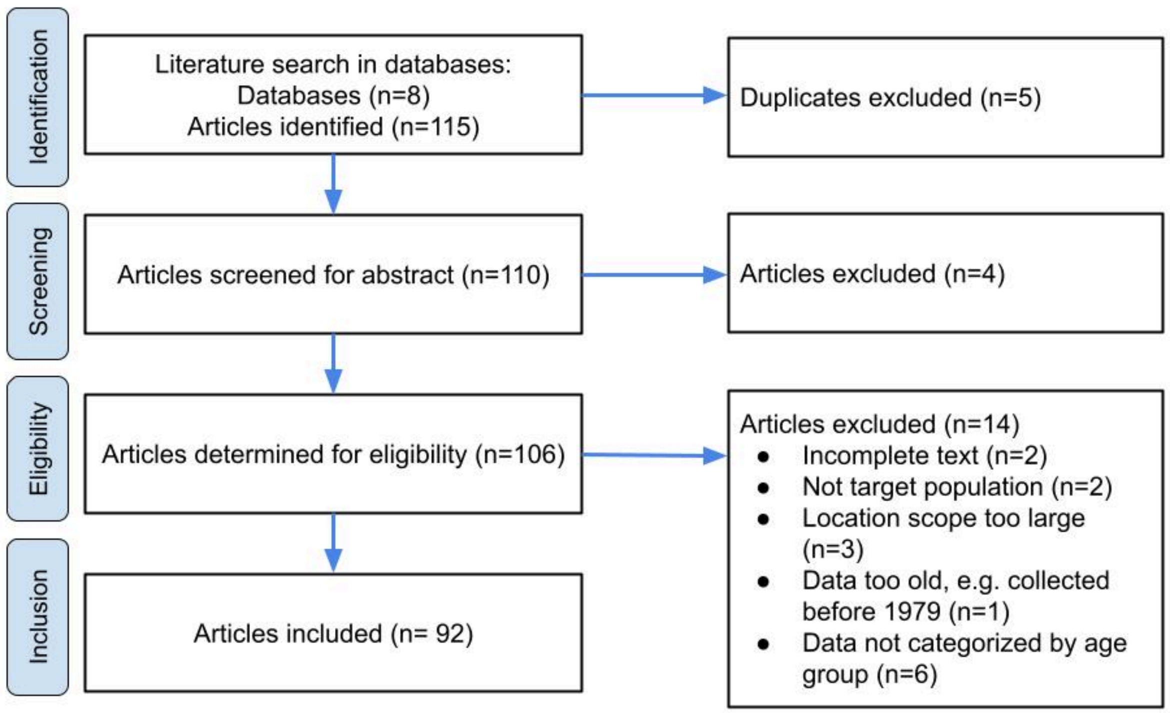


**Figure S1: Selection process of studies according to the PRISMA flow diagram.**

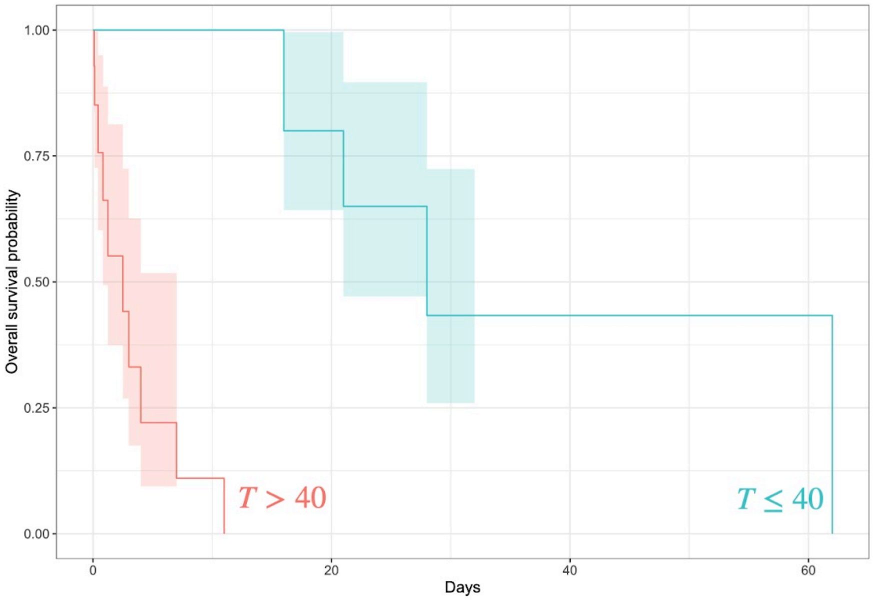


***Figure S2*****: Effect of temperature on oocyst survival.** Dubey et al. [10] evaluated success or failure of experimental infection of mice following oocyst exposure to different temperatures (35, 40, 45, 50 and 55 degrees for variable durations; n= 62, with 35 events, see Supplementary R Code). Kaplan Meier estimates indicate that above 40 degrees celsius, oocyst survival (measured as ability to infect susceptible mice) is considerably reduced.

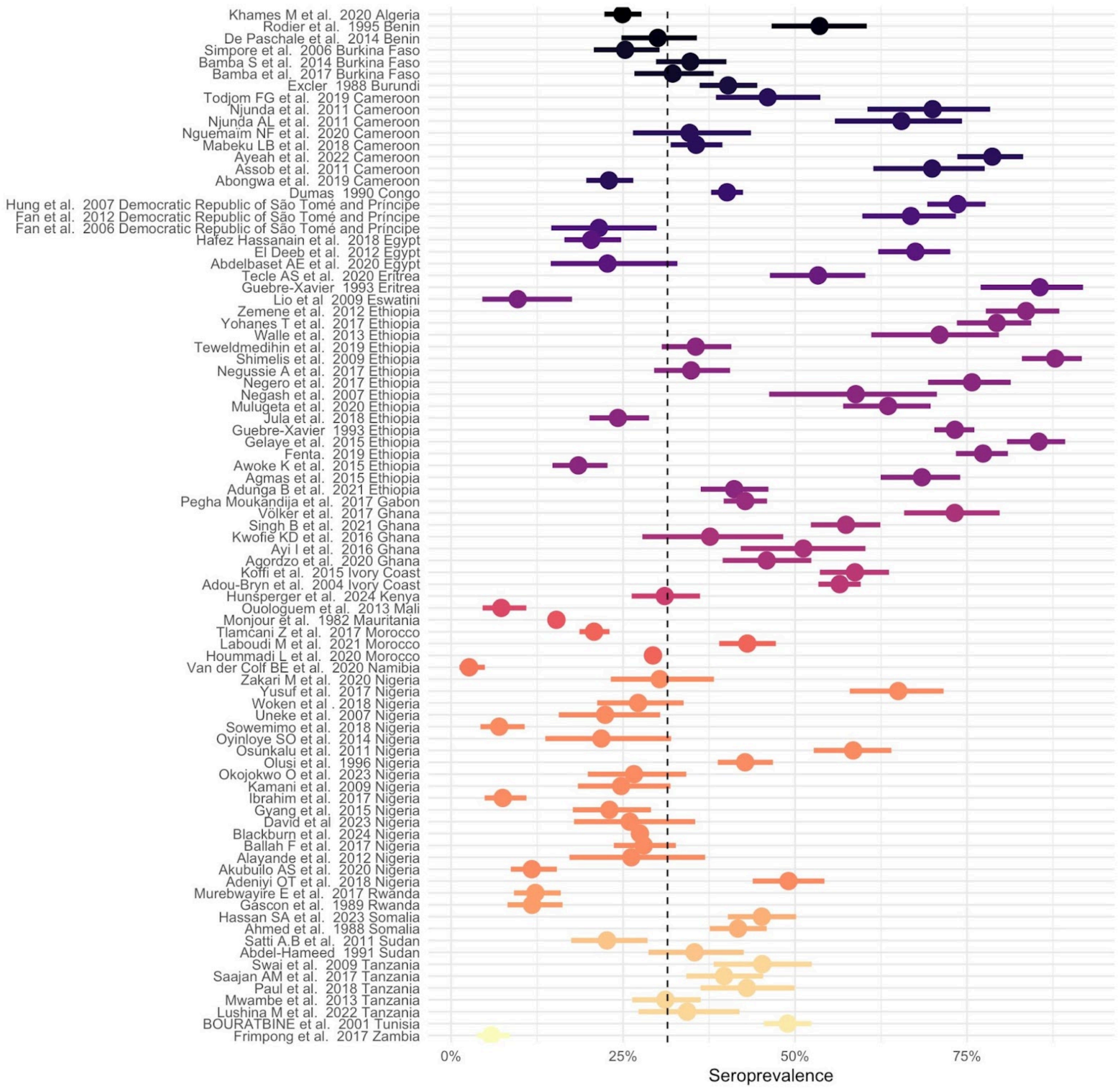


Figure S3: Forest plots of *Toxoplasma* seroprevalence in pregnant women in Africa between 1979 and 2025. Dashed line indicates pooled seroprevalence from 83 919 pregnant women across x countries (seroprevalence: 31.4%, 95% CI : 31%-32%)
